# Abnormal (hydroxy)prolines deuterium content redefines hydrogen chemical mass

**DOI:** 10.1101/2021.11.11.468216

**Authors:** Hassan Gharibi, Alexey Chernobrovkin, Gunilla Eriksson, Amir Ata Saei, Zena Timmons, Andrew C. Kitchener, Daniela Kalthoff, Kerstin Lidén, Alexander A. Makarov, Roman A. Zubarev

## Abstract

Analysing the δ2H in individual amino acids of proteins extracted from vertebrates, we unexpectedly found in some samples, notably bone collagen from seals, more than twice as much deuterium in proline and hydroxyproline residues than in seawater. This corresponds to at least four times higher δ2H than in any previously reported biogenic sample. We ruled out diet as a plausible mechanism for such anomalous enrichment. This finding puts into question the old adage that you are what you eat.

**SUMMARY:** The chemical mass of hydrogen is defined as an interval from the lowest to the highest content of deuterium ^2^H, hydrogen’s heavy stable isotope. Measurements of the deviations δ^2^H in the deuterium content from the standard (ocean water, δ^2^H = 0‰) are used to characterise biological samples, such as animal bone collagen. The results are often interpreted in terms of the trophic level and diet of the animal as well as prevailing climate during its lifetime. The majority of the published bone collagen δ^2^H data fall into a narrow δ^2^H range limited to ±100‰. Using novel analysis method, we unexpectedly found greatly higher δ^2^H values, up to 1500‰, in seal bone collagen. Such anomalous deuterium enrichment is detected only in two amino acid residues, proline and its derivative hydroxyproline, while other residues show much smaller δ^2^H values. Anomalously high δ^2^H values, albeit of lower magnitudes, are also found for these residues in other biological sources. This finding substantially expands the upper bound of the hydrogen chemical mass for biogenic sources. Since neither diet nor environment explain these mysteriously high enrichment levels amounting to more than twice deuterium content in sea water, our understanding of stable isotopes in nature, as well as the old adage “you are what you eat”, are put in question.

## Main Text

Deuterium ^2^H is a stable isotope of hydrogen present in nature at the level of 120-150 parts per million (ppm). Measurements of the deviations δ^2^H in deuterium content from standard (ocean water, δ^2^H = 0‰) are widely used in science for terrestrial ^1^ as well as extra-terrestrial ^2,3^ samples. An important biogenic sources for δ^2^H analysis is collagen, the most abundant protein in bones and skin ^4^. Collagen molecules form strong fibrils that can survive thousands, if not millions, of years, after an animal’s death ^5^ due to the protective bone matrix ^6–8^. Deuterium content together with ^13^C and ^15^N abundances are used to derive information on the habitat and diet of the animals, some long extinct, as well as the climate in their lifetime ^9–11^.

Most of the collagen δ^2^H measurements that have been reported to date in scientific literature fall into a limited δ^2^H range, usually ±100‰ (10%) ^12^. The highest published δ^2^H values, ∼200‰, are found in bone collagen of adult marine piscivores ^12,13^. In 2008, a record 299‰ enrichment in collagen from a seal bone was reported in the Quoygrew medieval burial site on the Orkney Islands, Scotland ^14^.

Here we found δ^2^H values in seal bone collagen that are several times higher than that, greatly surpassing the upper bound of hydrogen chemical mass recorded so far in any biogenic sample. Unlike the previous studies concerning bulk collagen, the reported extreme δ^2^H values are found only in two types of amino acid residues present in collagen, namely proline (Pro) and its post-translationally modified derivative hydroxyproline (Hyp). Given that Pro and Hyp compose >22% of all residues in mammalian collagen and that they are roughly twice as heavy as glycine, the most abundant collagen amino acid, our result is in broad agreement with the 2008 measurements performed on bulk protein. Yet our finding represents a dramatic extension of the range of deuterium content in natural substances, redefining the biogenic chemical mass for this important element (Fig. 1).

**Fig. 1.**
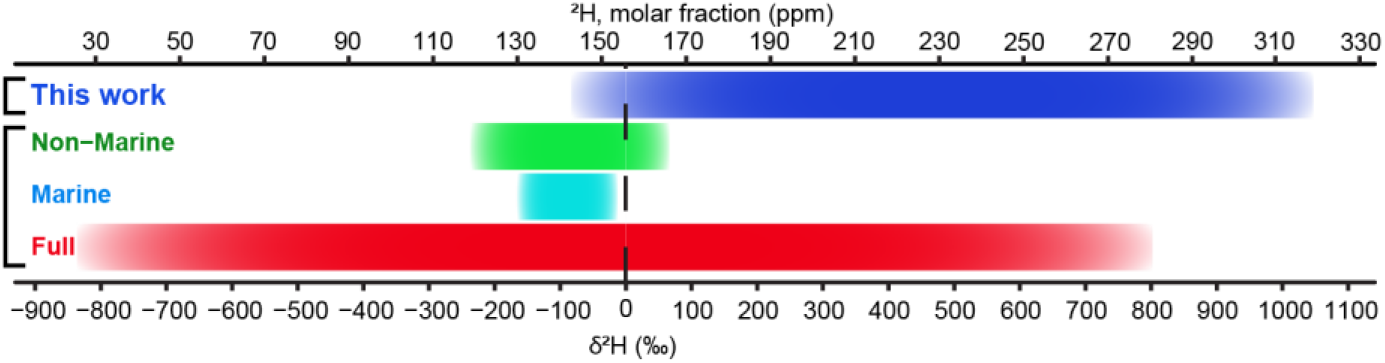
Deuterium range in nature according to IUPAC and its extension in this work

Isotopic ratio measurements with amino acid resolution in collagen were performed using the recent method of Fourier transform isotopic ratio mass spectrometry (FT IsoR MS, Fig. 2a). In short, collagen is extracted from animal bones as in conventional isotopic ratio mass spectrometry (IR MS) and then digested by trypsin as is common in proteomics. The obtained peptide mixture is analysed in a proteomics-type data-dependent experiment with liquid chromatography coupled with tandem mass spectrometry (LC-MS/MS), with the only difference being that, after the conventional MS/MS event generating sequencing information (which in FT IsoR MS is optional), an additional MS/MS event is introduced. This event uses a broad m/z window for precursor ion isolation, followed by hard gas-phase collision-induced dissociation, fragmenting collagen tryptic peptide ions down to immonium ions H_2_N^+^=CHR, where R represents the side chain of an amino acid residue. All aliphatic amino acids as well as aromatic ones give abundant immonium ions. For Pro and Hyp, the immonium ions are cyclic, C_4_H_8_N^+^ 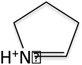 for Pro and C_4_H_8_NO^+^ 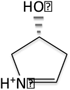 for Hyp. The mass resolution of 60,000 at m/z 200 of the Orbitrap mass analyser enables measurements of the “fine structure” M+1 peak components for ^2^H, ^13^C, and ^15^N isotopes (Fig. 2b). The specific isotopic ratio (e.g. ^2^H/^1^H) is obtained from the FT IsoR MS/MS spectrum as the abundance ratio of a specific component (^2^H-component in this communication) of the fine structure of an (M+1) isotopic peak and the abundance of the monoisotopic peak M in a given tandem mass spectrum, and dividing this ratio by the number of atoms of the corresponding element in the immonium ion (in case of hydrogen, seven for Pro and eight for Hyp ions). In an LC-MS/MS analysis of collagen peptides, several hundreds of individual isotopic measurements are obtained, forming a bell-shaped distribution (Fig. 2c). The most probable value of this distribution provides the final δ^2^H result. The δ^2^H variation between replicate analyses is ≤3‰ in case of abundant immonium ions, such as Pro, Hyp and Ile/Leu. To validate the FT IsoR MS method, we performed analysis of a proxy standard composed of free amino acids individually analysed at the bulk level by two certified IR MS laboratories (Extended Data Figure 1).

**Fig. 2.**
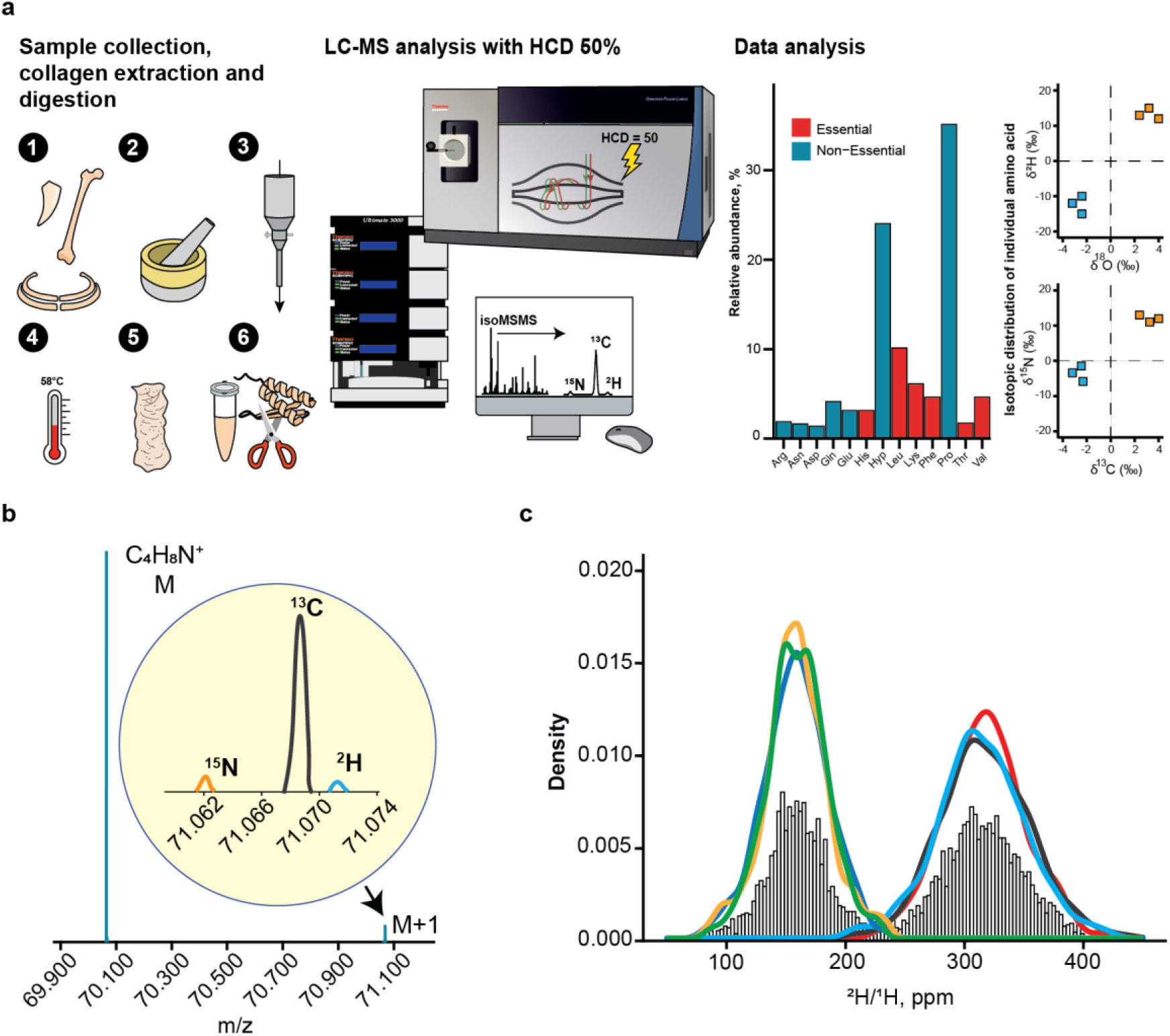
A. The workflow of FT isoR MS. 1: bone sampling; 2: bone homogenisation; 3: bone decalcification with HCl, 4: bone powder hydrolysis in diluted HCl at elevated temperature; 5: lyophilized collagen; 6: in-solution trypsin digestion, followed by LC-MS/MS and data processing. B. The monoisotopic peak M and the “fine structure” M+1 peak components for ^2^H, ^13^C, and ^15^N isotopes of a Pro immonium ion. C. Distributions of the ^2^H/^1^H ratios for Pro immonium ion obtained during a 1 h LC-MS/MS analysis of two distinct collagen samples (swan and grey seal) in three replicates each; the reported value is the mean of the average values in each replicate distribution.

The FT IsoR MS results for grey seal collagen in comparison with swan collagen (the latter was chosen as an interim standard) are shown in Fig. 3a. In Pro immonium ion 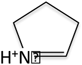, the nitrogen-bound hydrogen is labile, and thus the observed enrichment is diluted as 7/8 by the δ^2^H in the LC solvent, which was taken to have the ^2^H/^1^H ratio of 147 ppm. In Hyp immonium ions 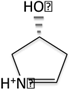, two hydrogen atoms are labile, and thus the dilution was 6/8. After correction was made for the dilution effect, the maximum Pro δ^2^H value was 1209‰, and for Hyp δ^2^H it was 1468‰. At the same time, for the Leu/Ile immonium ions a slight depletion (−6‰) was observed.

**Fig. 3.**
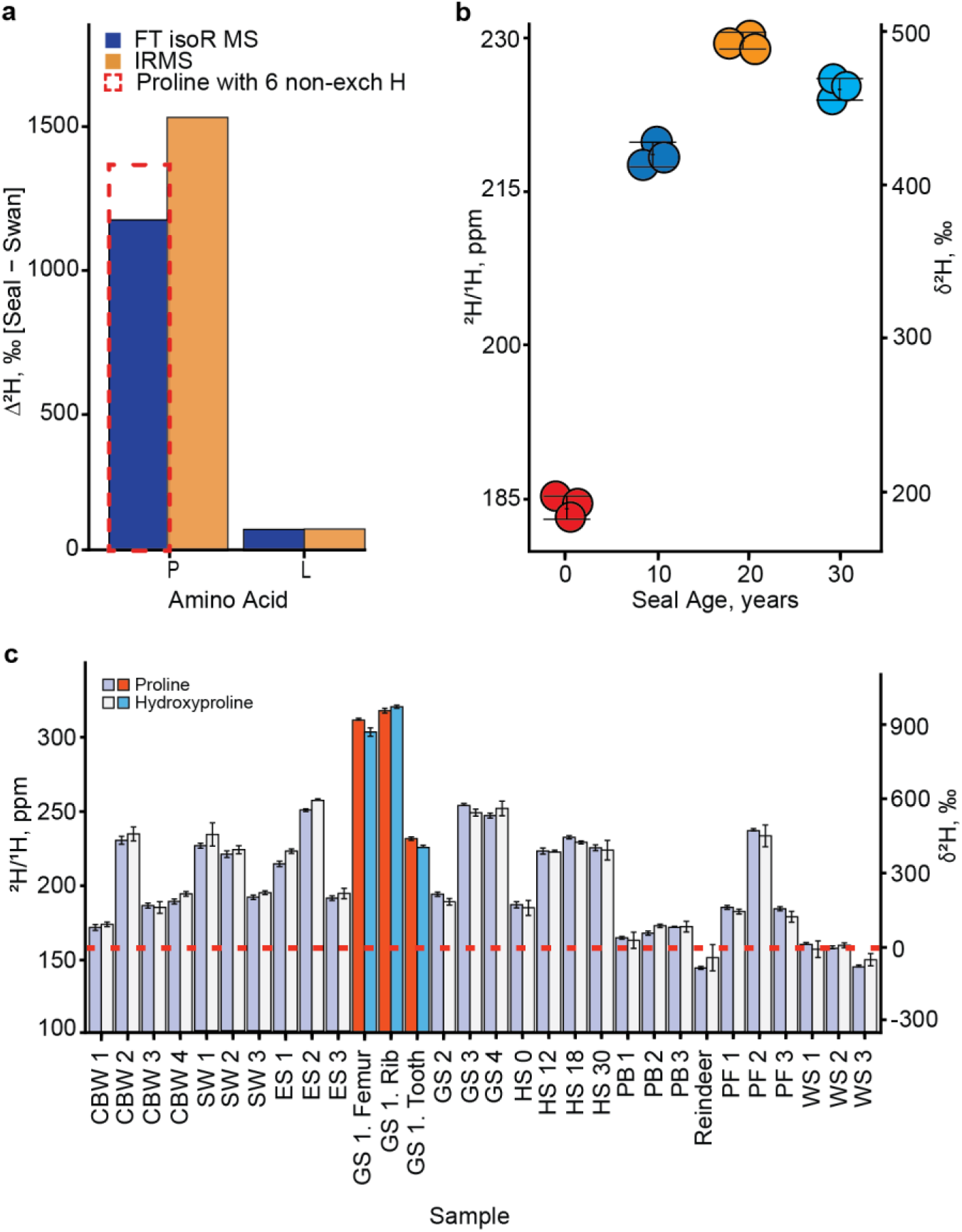
A. Bar plot shows the δ^2^H difference between seal and swan samples in IRMS and FT isoR MS (Dashed plot for different number of labile hydrogens) for proline (P) and leucine (L). B. Deuterium accumulation with age in 4 harbor seals; each sample is analyzed in 3 replicates. C. Deuterium content in Pro and Hyp in analyzed samples, CBW = Cuvier’s Beaked Whale, ES = Southern Elephant Seal, PF = Peregrine Falcon, GS = Grey Seal, HS = Habour Seal, PB = Polar Bear, SW = Sperm Whale, WS = Whooper Swan.

These extraordinary results were verified by amino acid-resolved isotope ratio MS, in which collagen is first hydrolysed to free amino acids that are then derivatized to become volatile, separated by gas chromatography and combusted, with the resultant H_2_ analysed by a magnetic sector mass spectrometer ^15^. The obtained δ^2^H value for Pro, 1700‰, corresponds to even higher enrichment than the FT IsoR MS value (Hyp was not analysed). Of the 12 amino acids analysed (Ala, Gly, Ser, Pro, Asp, Glu, Thr, Val, Leu, Ile, Phe and Lys), all but Gly showed enrichment in seal versus swan, though none as much as Pro (Extended Data Table 1). The second-most enriched amino acid, Thr, had a δ^2^H value >3 times lower than that of Pro.

We also analysed with FT IsoR MS bone collagen from the bones of various animals collected in different geographical areas (Extended Data Figure 2) dating from the Medieval age until modern times and samples with different biological ages to have a more comprehensive analysis (Fig. 3b,c). Many samples showed elevated deuterium levels in Pro and Hyp: e.g., the fastest flying bird, peregrine falcon (speeds up to 320 km/h ^16,17^) showed δ^2^H values up to 500‰ (Fig. 3c). Among different seal species, grey seals (*Halichoerus grypus*) from Scotland had the highest values (Fig. 3c). There seems to be a tendency for δ^2^H values to increase with biological age in seals, although data do not exclude saturation at puberty (Fig. 3b). At the same time, in all the analysed species Ile/Leu residues had unremarkable δ^2^H values near zero (Extended Data Table 2).

Finding an explanation for this phenomenon turned out to be problematic. Pro is a non-essential amino acid in mammals, and it can be synthesised from both glutamate and ornithine ^18^. In IRMS analysis, however, Glu and Lys (the closest analogue of ornithine) were the 5^th^ and 9^th^ most enriched residues out of 12 (Extended Data Table 1), which makes *de novo* biosynthesis of Pro an unlikely cause of deuterium enrichment.

Diet is also a doubtful explanation: in laboratory experiments on mice, δ^2^H in adult bone collagen was always below that of food ^19^. Also, δ^2^H Pro and Hyp values in polar bears that feed mostly on seals were only slightly above the reindeer and much below those of the seals (Fig. 3c).

Looking for other explanations, we considered radical reactions that can be initiated in proteins by mechanical stress ^20^. In structurally stressed peptides, the acidities of the alpha-proton are higher in prolyls than other amino acid residues abundant in collagen ^21^, and thus prolyl alpha-carbon is a likely site of hydrogen abstraction. And as the C-D bond is stronger than the C-H bond ^22^, deuterium atoms at alpha-carbons should be less likely abstracted and more readily attached to alpha-carbon radicals than hydrogen atoms. However, when bone collagen was ground in a ball mill containing water, no significant alteration in proline deuterium was detected at different deuterium contents in water.

Another hypothesis to check was whether deuterium enrichment could be caused by stress during the growing phase of the collagen-producing cells. To test this possibility, we grew human fibroblasts under different stress conditions (water shortage, food starvation, and temperature fluctuations) for six weeks. Neither of the tested stress conditions cause deuterium enrichment in Pro exceeding the media ^2^H content.

We further hypothesized that, since proline side chains easily make complexes with metal (II) ions, and such complexes, at elevated temperature, undergo hydrogen-deuterium exchange ^23,24^, the presence of these ions that stabilize collagen filaments in bones ^25,26^ can lead to deuterium accumulation in proline. To test this hypothesis, we incubated collagen peptides with Zn^2+^, Cu^2+^, Mn^2+^ and Mg^2+^ salts at 90 °C for 72 h in water containing normal as well as elevated deuterium content. No enrichment in proline compared to δ^2^H in water was observed.

Remaining a mystery, the biochemical pathway for deuterium enrichment in Pro and Hyp residues of proteins calls for further investigation of this intriguing phenomenon.

## Acknowledgements

This work was supported by the Swedish research council (grant 2017-04303 to RAZ). Amino acid-resolved IR MS analysis was performed at the Department of Earth and Planetary Sciences, University of California Riverside (UCR), USA. Kaycee Morra, Seth Newsome and Marilyn L. Fogel of UCR are acknowledged for helpful discussions.

## Author Contributions

Hassan Gharibi and Alexey Chernobrovkin has contributed equally to this paper.

## Supporting Information available

Details of the mass spectrometric analyses used to generate the data, as well as Excel files containing all information are available upon request.

## Additional Information

